## Supplementary Figures for "An evolutionary functional genomics approach identifies novel candidate regions involved in isoniazid resistance in *Mycobacterium tuberculosis*"

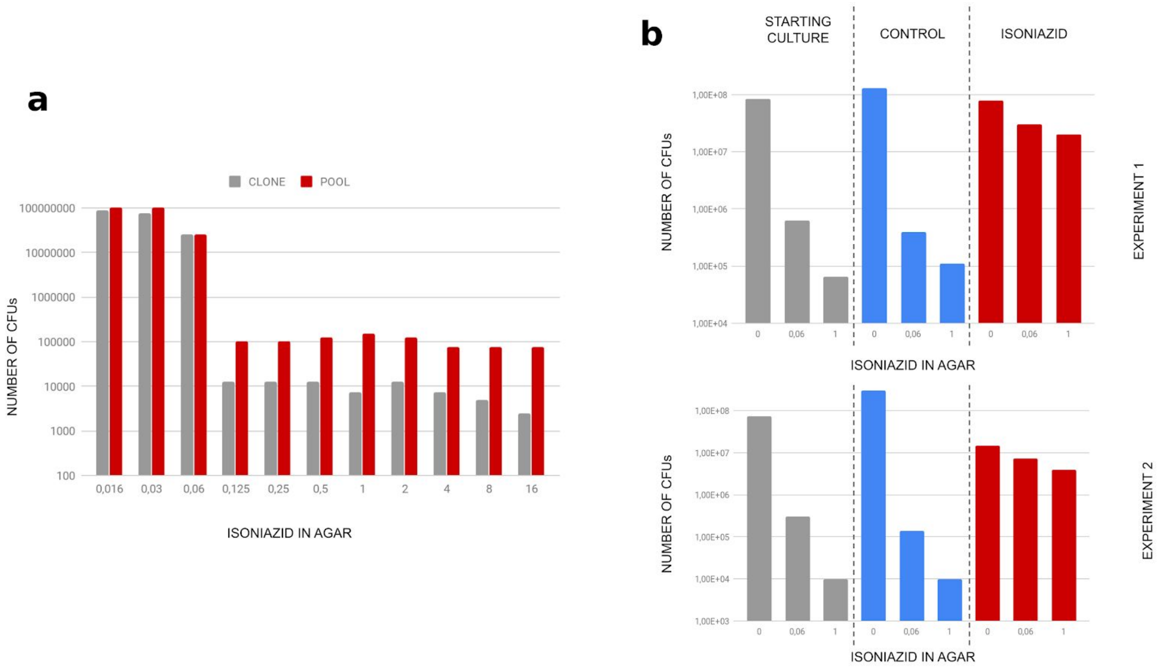


**Supplementary Figure 1: a**) The pool contains 10 times more isoniazid resistant mutants than its parental clone. **b**) Throughout the experiment, the frequency of isoniazid resistant mutants increased specifically in populations treated with the antibiotic.

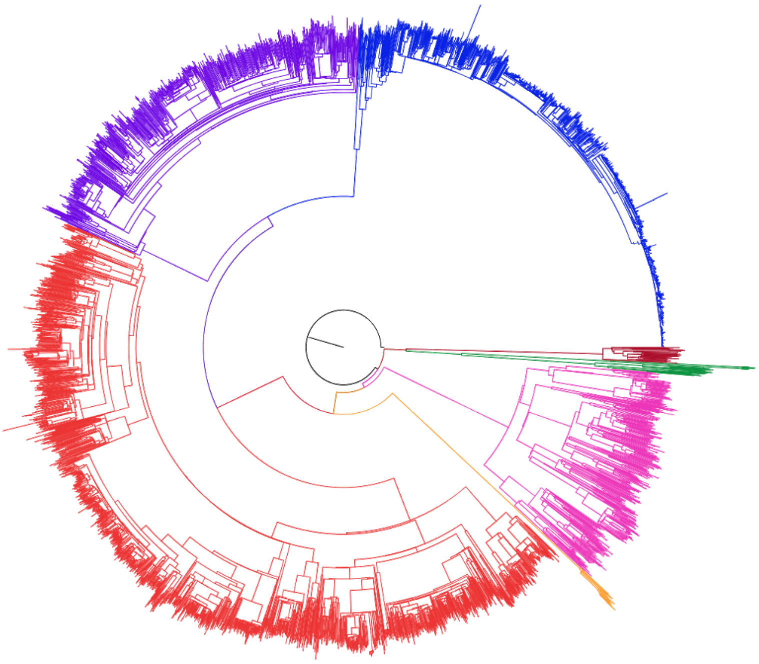

**Supplementary Figure 2:** Global phylogeny generated with a collated dataset from various sources. Interactive phylogeny can be found at https://itol.embl.de/tree/161111218247308411581608730

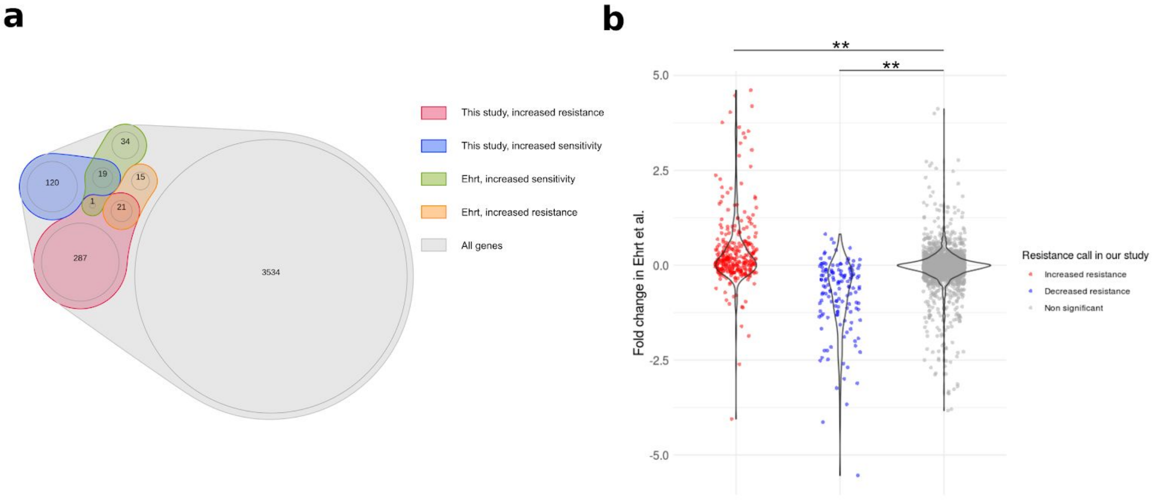


**Supplementary Figure 3: a**) Coincidence in resistance-altering features detected by functional genomics between our study and that of Ehrt *et al.* **b**) Our resistance-altering features showed similar phenotypes in Xu, W. *et al.* Chemical Genetic Interaction Profiling Reveals Determinants of Intrinsic Antibiotic Resistance in Mycobacterium tuberculosis. *Antimicrob. Agents Chemother.* **61**, (2017).
